# Ion Gradient-driven Bifurcations of a Multi-Scale Neuronal Model

**DOI:** 10.1101/2022.10.01.510461

**Authors:** Anthony G. Chesebro, Lilianne R. Mujica-Parodi, Corey Weistuch

## Abstract

Metabolic limitations within the brain frequently arise in the context of aging and disease. As the largest consumers of energy within the brain, ion pumps that maintain the neuronal membrane potential are the most affected when energy supply becomes limited. To characterize the effects of such limitations, we analyze the ion gradients present in the Larter-Breakspear neural mass model. We show the existence and locations of Neimark-Sacker and period-doubling bifurcations in the sodium, calcium, and potassium reversal potentials, and demonstrate that these bifurcations form physiologically relevant bounds of ion gradient variability. Within these bounds, we show how depolarization of the gradients will cause decreased neural activity. We also show that the depolarization of ion gradients decreases inter-regional coherence, causing a shift in the critical point at which the coupling occurs and thereby inducing loss of synchrony between regions. In this way, we show that the Larter-Breakspear model captures ion gradient variability present at the microscale level and propagates these changes to the macroscale effects congruent with those observed in human neuroimaging studies.

## 1. Introduction

Although by mass the brain is relatively small, it consumes roughly 20% of the body’s energy [1]. The majority of this energy is expended by neuronal ion pumps maintaining the ion gradients necessary for action potentials [1, 2]. Thus, the brain is particularly affected by metabolic energy deficits [3, 4, 5]. Recent work has pointed to metabolic energy deficits as causing brain network instability [5] and steepening cognitive decline in the context of aging [4]. Conversely, providing access to energy acutely (e.g., via ketone delivery) can have a stabilizing effect, underlining the crucial role of energy regulation in preserving homeostasis of neuronal computation [5, 6].

As the largest consumers of energy in the brain, ion pumps are one of the cellular functions most susceptible to metabolic deficits [1, 2, 3, 7, 8]. Ion pumps, and the ion gradients they generate across the cell membrane, are subject to tight regulation [9] and provide signal feedback to metabolic centers within the cell to ensure energy production meets demand [1]. Incorporation of ion gradients in computational neuron models has been essential, as they drive the ability to fire action potentials [10, 11]; however, the perturbation of ion gradients caused by metabolic constraints has only been a feature of more recent single neuron models [2, 11].

Single neuron models have proven invaluable in understanding the effects of ion gradient depletion, but they remain challenging to scale to whole-brain dynamical systems. The Larter-Breakspear neural mass model [12, 13], a multi-network extension of the Morris-Lecar equations [10], provides a computational unit that estimates the mean firing rates and membrane voltages of many neurons simultaneously. This makes it computationally feasible to produce a whole-brain simulation comprising many of these regions [14]. Crucially, the Larter-Breakspear model also retains explicit ion gradient dynamics, allowing manipulation of the gradients to simulate metabolic constraints without requiring single neuron scale computation.

In this work, we present a bifurcation analysis of the ion gradients for sodium, calcium, and potassium in the Larter-Breakspear model. Bifurcation analyses have been used to analyze the oscillations of many chaotic systems [15, 16], encompassing both neural models [17, 18, 19, 20, 21] and other biological systems [22, 23]. As the Larter-Breakspear model is a chaotic system [12, 13], bifurcation analysis allows us to rigorously analyze the ion gradients for critical points and thereby establish limits of physiologically relevant variability for future simulations. Having derived these boundaries, we then demonstrate how the mean membrane voltage of the neural mass is sensitive to each gradient. We show lengthened refractory periods (in the case of sodium/calcium hyperpolarization or potassium depolarization), increased degenerate oscillations (in the case of sodium/calcium depolarization), and increased oscillation frequency (in the case of potassium hyperpolarization). Finally, we show how the variability in the ion gradients alters the coupling between two different regions, altering the critical point where regions transition from chaotic to coupled oscillations, with depolarization of ion gradients causing reduced synchrony.

## 2. Model motivation and design

### 2.1. Neurobiology of the Larter-Breakspear model

The Larter-Breakspear model is constructed to extend ion dynamics from single neuron modeling to a neural network, allowing for greater scalability [13, 24]. There are several kinds of ion pumps and channels within a neuron (Fig. 1a). In this model, we are most concerned with those that require energy to function, such as the Na^+^/K^+^ ATP-ase [8], which generates a net positive sodium potential *V*_Na_ and a net negative potassium potential *V*_K_, and the plasma membrane Ca^2+^ ATP-ase [25], which generates a net positive calcium potential *V*_Ca_. There are also non-energetically active ion leak channels that allow several ion species to travel down their electrochemical gradients; these are considered in bulk as the potential *V*_L_.

**Figure 1:**
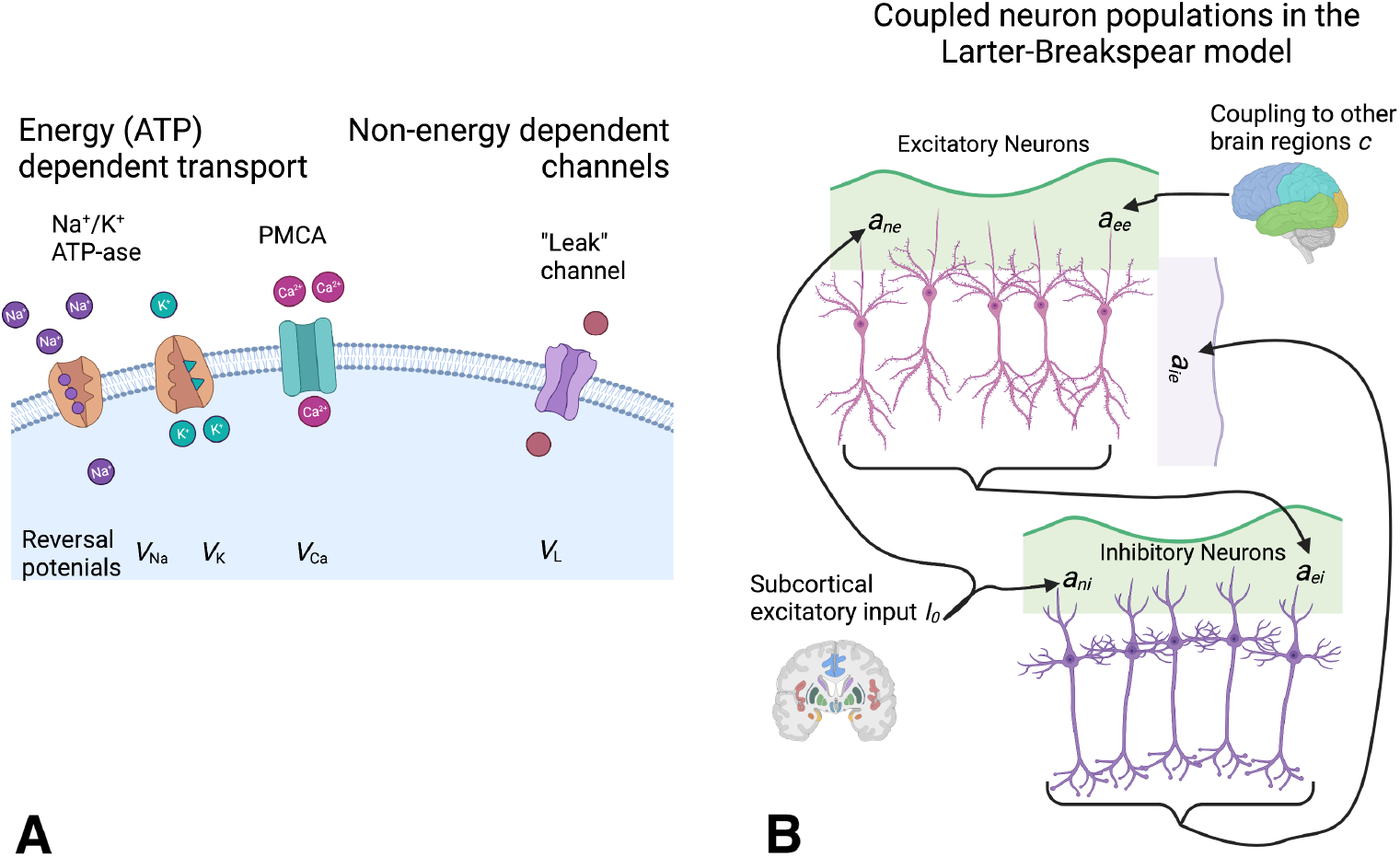
Microscale neurobiological components are used to construct the Larter-Breakspear model. A. Illustration of the ion transporters and channels used in the Larter-Breakspear model. Each of the relevant reversal potentials (*V*_Na_, *V*_K_, *V*_Ca_ and *V*_L_) are used to build two types of neuron populations: excitatory (pink) and inhibitory (purple). B. Each neural mass in the model is comprised of a an excitatory subnetwork and an inhibitory subnetwork, coupled together by various weights as shown (*a_ee_*, *a_ei_*, and *a_ie_*). Subcortical input *I*_0_ is received by both populations, while coupling between neural masses is done via the excitatory pathway only.

Utilizing these ion transporters as building blocks, the Larter-Breakspear model connects two subnetworks of neurons together to form a single mass: a population of excitatory neurons with mean membrane potential *V* and a population of inhibitory neurons with mean membrane potential *Z*. The excitatory subnetwork feeds back on itself (*a_ee_*) and projects to the inhibitory subnetwork (*a_ei_*), while also receiving feedback from the inhibitory neurons (*a_ie_*). Subcortical input *I*_0_ is received by both the excitatory (*a_ne_*) and inhibitory (*a_ni_*) subnetworks. Finally, two regions (each comprised of their own excitatory and inhibitory subnetworks) can be coupled together (*c*) through interactions between their excitatory subnetworks.

### 2.2. Larter-Breakspear model - single neural mass

Using the previous neurobiology to inform the model, the Larter-Breakspear equations are constructed as a system of three variables: mean excitatory membrane voltage *V*(*t*), mean inhibitory membrane voltage *Z*(*t*), and the proportion of potassium channels open at a given time *W*(*t*). Note that while *V*(*t*), *Z*(*t*), and *W*(*t*) are all time-dependent, we omit this dependence in the following equations for ease of reading. Given this understanding, the Larter-Breakspear model is defined as:

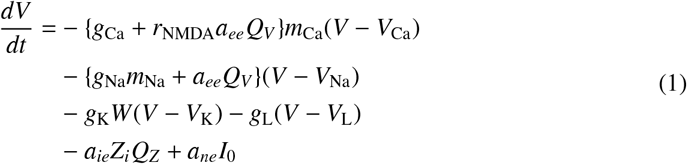

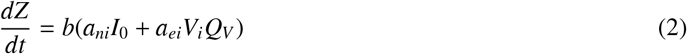

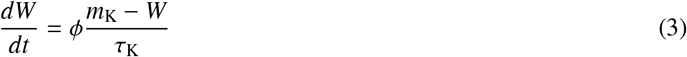

In these equations, *Q_V_* and *Q_Z_* are the mean firing rates for excitatory and inhibitory cell populations, respectively. These are computed as

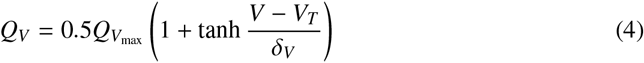

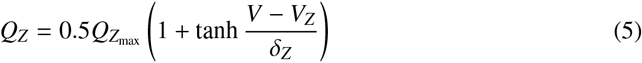

The individual ion channel gating functions (*m*_Na_, *m*_K_ and *m*_Ca_) take the form

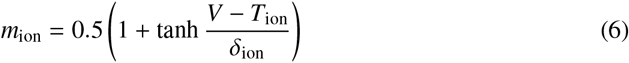

where *m*_ion_ is the fraction of voltage-dependent channels open at any given time. Default values and descriptions for all constants in these equations are given in Table 1. Note that parameter values are unit-less to scale to a reasonable modeling range (i.e., *V*, *Q* ∈ (−1, 1) and *W* ∈ (0, 1)), and the integration time step *dt* is in milliseconds.

**Table 1:** Default constant values in the Larter-Breakspear model. In Sections 3 and 4, we explore changes in the first three ion gradients, altering the reversal potentials *V*_Na_, *V*_K_, and *V*_Ca_. *The selection of *δ*_*V*,*Z*_ has been the subject of much prior work; see Appendix A for a more detailed consideration. **This is the coupling constant needed to achieve marginal stability in a 2-ROI system. See Section 4 for a more complete consideration of coupling constant range.

| Parameter | Description | Default Value |
| --- | --- | --- |
| **Nernst potentials** |  |  |
| $V_{Na}$ | Na<sup>+</sup> reversal potential | 0.53 |
| $V_K$ | K<sup>+</sup> reversal potential | -0.7 |
| $V_{Ca}$ | Ca<sup>2+</sup> reversal potential | 1.0 |
| $V_L$ | Leak channels reversal potential | -0.5 |
| **Channel conductances** |  |  |
| $g_{Na}$ | Na<sup>+</sup> conductance | 6.70 |
| $g_K$ | K<sup>+</sup> conductance | 2.00 |
| $g_{Ca}$ | Ca<sup>2+</sup> conductance | 1.00 |
| $g_L$ | Leak channels conductance | -0.50 |
| **Channel voltage thresholds** |  |  |
| $T_{Na}$ | Na<sup>+</sup> channel threshold | 0.30 |
| $T_K$ | K<sup>+</sup> channel threshold | 0.00 |
| $T_{Ca}$ | Ca<sup>2+</sup> channel threshold | -0.01 |
| **Channel voltage threshold variances** |  |  |
| $\delta_{Na}$ | Na<sup>+</sup> channel threshold variance | 0.15 |
| $\delta_K$ | K<sup>+</sup> channel threshold variance | 0.30 |
| $\delta_{Ca}$ | Ca<sup>2+</sup> channel threshold variance | 0.15 |
| **Excitatory & inhibitory population parameters** |  |  |
| $V_T$ | Excitatory neuron threshold voltage | 0.00 |
| $Z_T$ | Inhibitory neuron threshold voltage | 0.00 |
| $\delta_{V,Z}$ | Variance of excitatory/inhibitory thresholds | 0.66* |
| $Q_{V_{max}}$ | Excitatory population maximum firing rate | 1.00 |
| $Q_{Z_{max}}$ | Inhibitory population maximum firing rate | 1.00 |
| $a_{ee}$ | Excitatory → excitatory synaptic strength | 0.36 |
| $a_{ei}$ | Excitatory → inhibitory synaptic strength | 2.00 |
| $a_{ie}$ | Inhibitory → excitatory synaptic strength | 2.00 |
| $a_{ne}$ | Non-specific → excitatory synaptic strength | 1.00 |
| $a_{ni}$ | Non-specific → inhibitory synaptic strength | 0.40 |
| **Other parameters** |  |  |
| $I_0$ | Subcortical excitatory input | 0.30 |
| $b$ | Time scaling factor | 0.10 |
| $\phi$ | Temperature scaling factor | 0.70 |
| $\tau_K$ | K<sup>+</sup> relaxation time | 1.00 |
| $r_{NMDA}$ | NMDA/AMPA receptor ratio | 0.25 |
| $c$ | ROI-to-ROI coupling constant | 0.1** |

We note that the three ions of interest are modeled in three different manners. Sodium serves as the dominant shape determinant of the neural mass spiking activity as it has the highest net positive conductance coupled to its ion channel gating function. Calcium serves as a secondary support of the spiking activity, providing some of the amplitude to the spiking activity. However, because the calcium gating function is also coupled to the excitatory population firing rate and the ratio of NMDA/AMPA receptors, calcium more importantly provides feedback to the neural mass firing rate. Finally, due to its unique modeling as a separate differential equation, potassium plays a unique role in determining the frequency of spiking activity (at larger reversal potentials) and the duration of the refractory period (at smaller reversal potentials). As a consequence of this extra modeling step, potassium also has a different ion gradient landscape than sodium and calcium (discussed below).

### 2.3. Larter-Breakspear model - coupled neural masses

Equations 1-3 describe a single neural mass comprised of two subnetworks. Coupling between pairs of neural masses (*i* and *j*) can also be achieved through connection terms:

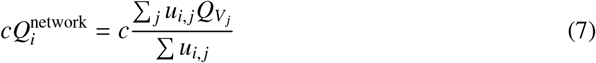

Here, *c* is the coupling constant, 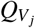 is the mean excitatory firing rate of region *j*, and *u_i, j_* is the strength of connection between regions *i* and *j*. The associated multi-regional Larter-Breakspear equations are then given by:

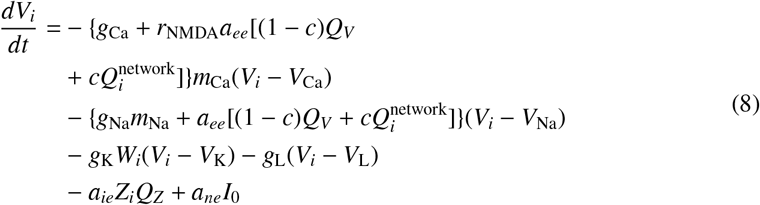

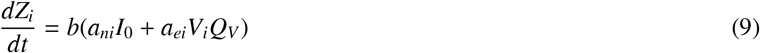

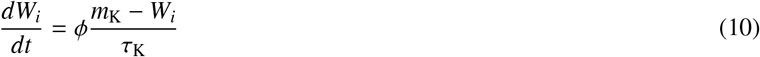

In this work, we use Equations 1-3 to probe the activity of individual neural masses (Section 3) and Equations 7-10 to explore the dynamics of two coupled regions (Section 4).

## 3. Codimension 1 bifurcations in the Larter-Breakspear model

### 3.1. Neimark-Sacker and period-doubling bifurcations

In this analysis, we show the existence and the significance of codimension 1 bifurcations (bifurcations that occur as a single parameter is varied) that exist in the sodium, calcium and potassium ion gradients in the Larter-Breakspear model. We specifically consider the Neimark-Sacker (or torus) bifurcation and the period-doubling (or flip) bifurcation. Torus bifurcations arise when a fixed point of the system (here the ion reversal potential) changes stability, and are characteristic of quasi-periodic oscillations of chaotic systems [21]. Flip bifurcations give rise to a dramatic shift in the period of the system as they are crossed (hence the term period-doubling) and form a plausible boundary of the ion gradients considered here. Proceeding beyond the flip bifurcation in an ion gradient typically causes a dramatic loss of frequency, with corresponding lengthening of refractory period that is no longer physiological. In spiking neuron models, the space between a torus and a flip bifurcation has been shown to exhibit rapidly changing frequency and quasi-periodicity, both features retained in the spaces we demonstrate below [21, 26].

Although the detection of codimension 1 bifurcations is possible analytically, in complex systems such as the Larter-Breakspear model it is often more feasible to establish their locations through numerical analysis [16]. To find the torus and flip bifurcations present in the Larter-Breakspear model, we employed the MatCont software package available for MATLAB [27]. This software allows for numerical detection of critical points, as well as the detection of limit cycle families and bifurcations. MatCont employs an implementation of the Moore-Penrose continuation near critical points; for further details on the computation of limits and detection of bifurcation points see Dhooge et al. [27] and Kuznetsov [28].

### 3.2. Bifurcations in the sodium ion gradient

We first consider the bifurcations arising as the sodium reversal potential *V*_Na_ is varied. As previously mentioned, this reversal potential is determined by the ion gradient maintained by the energy-dependent Na^+^/K^+^ ATP-ase in the neuron cell membrane, with a net positive charge induced by a surplus of sodium ions outside the cell. Physiologically, this reversal potential is 68mV [2], which corresponds to a resting reversal potential of 0.53 (hereafter referred to as baseline) in the Larter-Breakspear re-scaled parameters.

Initiating a search for critical points near the equilibrium point of the system, we find that there is a Hopf point in the sodium gradient at *V*_Na_ ≈ 0.2432, significantly depolarized relative to the baseline reversal potential. (For consideration of the other points discovered using this method, see Appendix B). Using Moore-Penrose continuation to detect the family of periodic orbits originating at the Hopf point, we gradually increase the value of *V*_Na_, producing the family of orbits shown in Fig. 2a. Highlighted in red are the two limit cycles corresponding to the torus and flip bifurcations (labeled NS and PD, respectively). The torus bifurcation occurs at *V*_Na_ ≈ 0.401, while the flip bifurcation occurs at *V*_Na_ ≈ 0.603.

**Figure 2:**
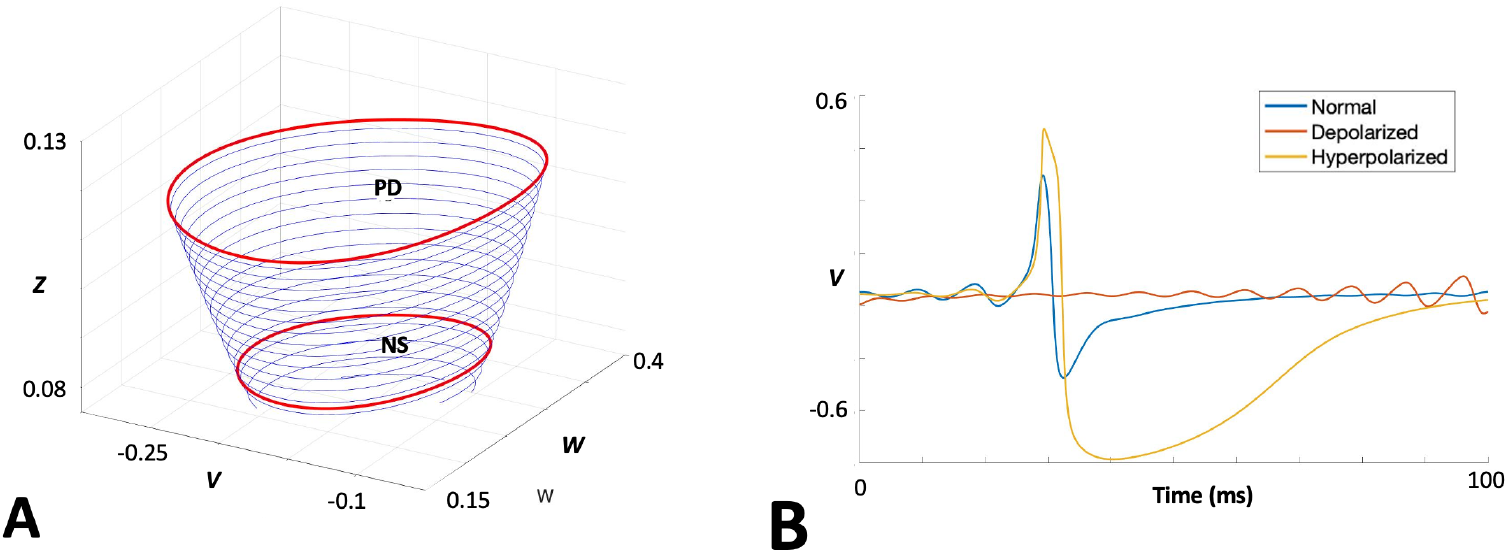
Torus and flip bifurcations in the sodium gradient alter refractory period and neuronal activity spiking. A. Family of limit cycles originating at the sodium gradient Hopf point (*V*_Na_ ≈ 0.24). The limit cycles corresponding to the Neimark-Sacker bifurcation (labeled NS) at *V*_Na_ ≈ 0.40 and the period-doubling bifurcation (labeled PD) at *V*_Na_ ≈ 0.60 are highlighted in red. B. Mean excitatory voltage *V* waveforms under three conditions: baseline reversal potential (*V*_Na_ = 0.53), depolarized just over the torus bifurcation (*V*_Na_ = 0.38), and hyperpolarized just over the flip bifurcation (*V*_Na_ = 0.68).

Hypothesizing that the space between these bifurcations is the physiologically relevant range of the sodium ion gradient, we probe the dynamic response of the neural mass to variations just beyond each bifurcation and analyze the change in dynamics of the mean excitatory membrane potential *V* (Fig. 2b). Hyperpolarization of *V*_Na_ just beyond the flip bifurcation leads to peaks in *V* of greater amplitude, a significantly prolonged recovery time, and decreased oscillatory activity immediately prior to the peaks. In stark contrast, depolarization of *V*_Na_ just below the torus bifurcation leads to a complete loss of sharp peaks, with only small, noisy oscillations occurring sporadically.

### 3.3. Bifurcations in the calcium ion gradient

Next, we consider the bifurcations in the calcium reversal potential *V*_Ca_. The resting reversal potential of calcium is roughly 140mV [11], giving a *V*_Ca_ = 1.0 at rest in the Larter-Breakspear model. Similar to the sodium results, when initiating a critical point scan near the equilibrium point we find a Hopf point at *V*_Ca_ ≈ 0.9098 (see Appendix B for other critical points). Detecting the family of periodic orbits originating from this point (Fig. 3a), we observe a torus bifurcation at *V*_Ca_ ≈ 0.959 and a flip bifurcation at *V*_Ca_ ≈ 1.024 (labeled NS and PD, respectively). Notably, this dynamic range is significantly smaller than that of *V*_Na_.

**Figure 3:**
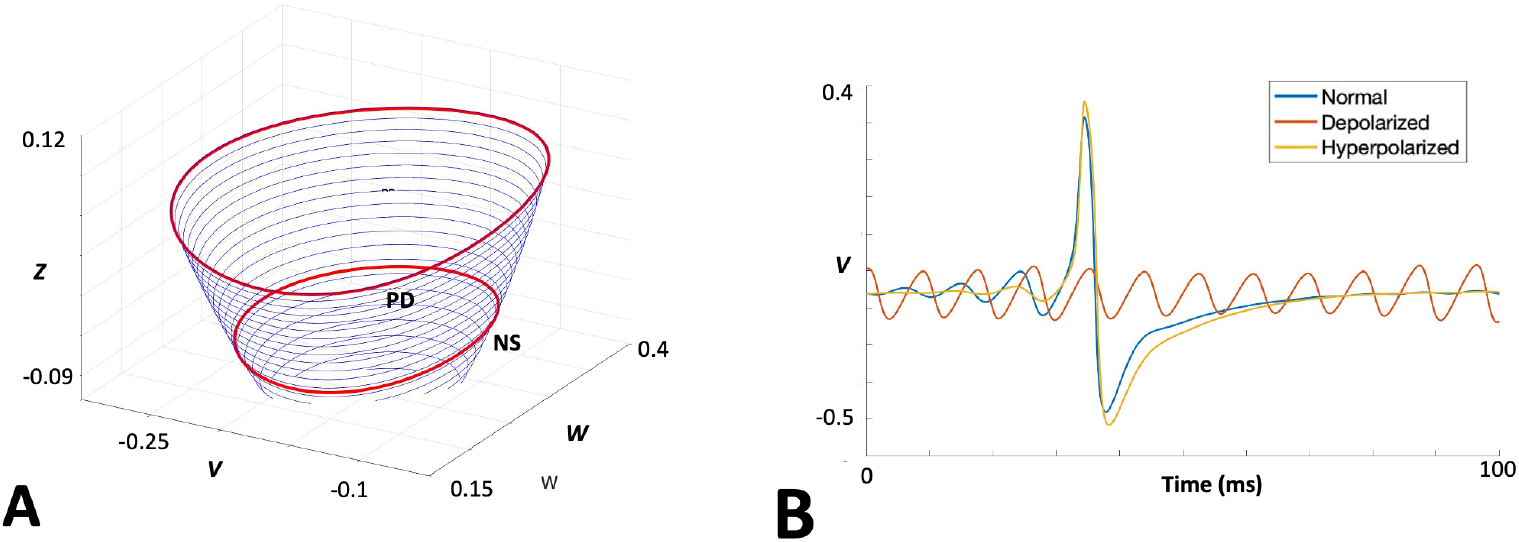
Torus and flip bifurcations in the calcium gradient alter neuronal activity spiking with minimal effect on refractory period. A. Family of limit cycles originating at the calcium graident Hopf point (*V*_Ca_ ≈ 0.91). The limit cycles corresponding to the Neimark-Sacker bifurcation (labeled NS) at *V*_Ca_ ≈ 0.96 and the period-doubling bifurcation (labeled PD) at *V*_Ca_ ≈ 1.02 are highlighted in red. B. Mean excitatory voltage *V* waveforms under three conditions: baseline reversal potential (*V*_Ca_ = 1.0), depolarized just over the torus bifurcation (*V*_Ca_ = 0.949), and hyperpolarized just over the flip bifurcation (*V*_Ca_ = 1.084).

Again taking the space between these two bifurcations as the range of physiological interest, we probed the dynamic response of *V* in response to crossing these bifurcations (Fig. 3b). We observe that hyperpolarization by crossing the flip bifurcation in the *V*_Ca_ gradient produces a markedly milder effect on the peak activity, with only a slight increase in the refractory period compared to a similar perturbation in *V*_Na_. However, we still observe a marked change in *V* dynamics crossing the torus bifurcation (depolarization *V*_Ca_), with the spikes subsiding into constant, high frequency, low amplitude oscillatory activity.

### 3.4. Bifurcation in the potassium ion gradient

Since potassium dynamics are computed with a separate differential equation in the Larter-Breakspear model, the same numerical analyses performed in the cases of sodium and potassium fails to find a Hopf point in the gradient, discovering only the other critical points (discussed further in Appendix B). However, we know from analytical observations that the change in dynamics is too steep to be continuous. The issue in discovering the critical point arises from the fact that the landscape of *W* is too smooth for the numerical computation to successfully identify a Hopf point. To make the landscape slightly more rough to assist the computation, we reduced the K^+^ relaxation time to τ_K_ = 0.9, which allowed us to proceed with the numerical discovery. The resting reversal potential of potassium is roughly −80mV [2], which corresponds to *V*_K_ = −0.7 in the Larter-Breakspear system. In our slightly modified regime, a Hopf point is present at *V*_K_ ≈ −1.102. Detecting the family of periodic orbits originating from this point (Fig. 4a), we observe only a single flip bifurcation at *V*_k_ ≈ −0.610 (labeled PD) - and notably an absence of any torus bifurcation in the hyperpolarizing gradient direction.

**Figure 4:**
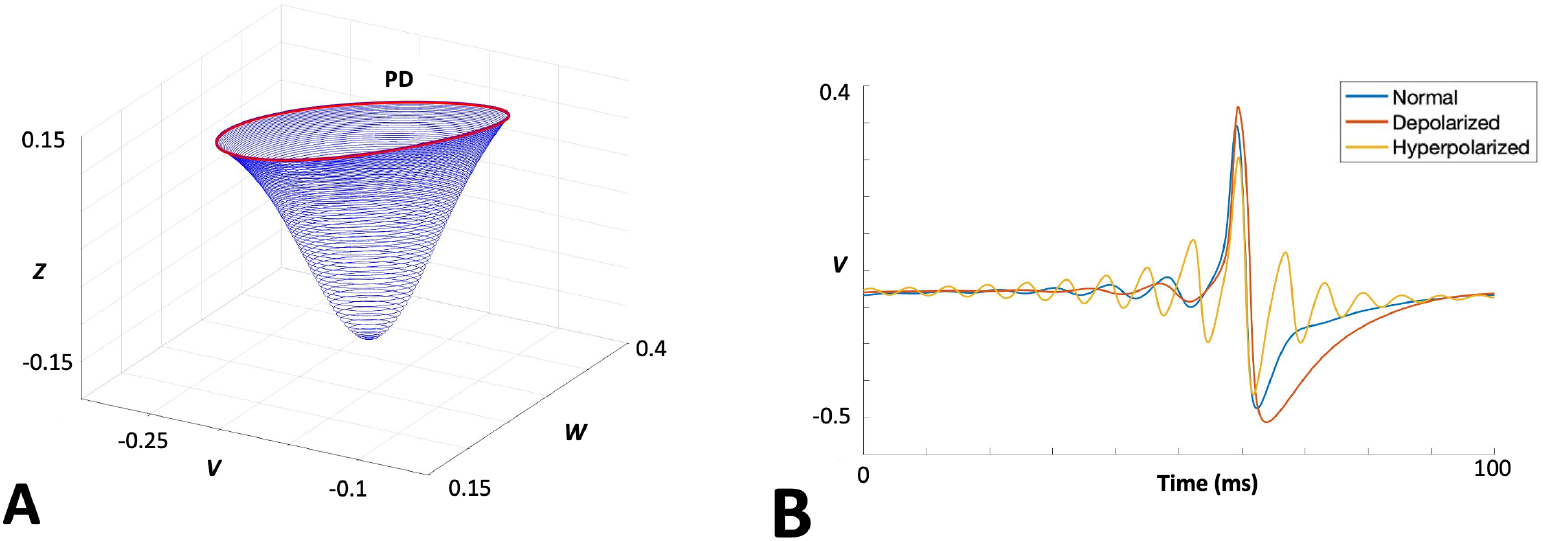
Flip bifurcation in the potassium ion gradient lengthens refractory period. A. Family of limit cycles originating at the potassium gradient Hopf point (*V*_K_ ≈ −1.102). Unlike the sodium and calcium gradients, there is no Neimark-Sacker bifurcation between the Hopf point and the period-doubling bifurcation, which here occurs at *V*_K_ ≈ −0.61 (shown in red and labeled PD). B. Mean excitatory voltage *V* waveforms under three conditions: baseline reversal potential (*V*_K_ = −0.7), depolarized just over the torus bifurcation (*V*_K_ = −0.59), and significantly hyperpolarized (*V*_K_ = 1.084).

Since there is no direct range of physiological values lying between two bifurcations to choose, we probed the dynamics of *V* at the resting reversal potential level, depolarized just over the flip bifurcation, and hyperpolarized by an arbitrary amount (but still below the limit point). We observe (Fig 4b) that crossing the flip bifurcation induces a significantly delayed refractory period, similar to what is observed in hyperpolarization of sodium. Hyperpolarizing *V*_K_ leads to a shortened refractory period, accompanied by significantly increased oscillation frequency without a distortion of spike shape.

## 4. Synchronization in varying ion regimes

To explore the relevance of these ion gradient bifurcation points in networks of coupled neural masses, we set up a system of two connected ROIs using Equations 7-10 above. Given prior work on the effects of metabolic constraints on brain network stability [4, 5], we hypothesized that depolarization of the membrane potentials - towards the torus bifurcation in the sodium and calcium gradients, and towards the flip bifurcation in the potassium gradient - would lead to an observable decrease in coupling between regions. To test this, we chose a coupling constant *c* near the zone of marginal stability (i.e., where the coupled system will alternate between chaotic and synchronized activity; see Breakspear et al. [13]) and varied the ion gradients across the physiologically relevant ranges identified in Section 3. We chose the covariance between the excitatory membrane voltages *V*_1_(*t*) and *V*_2_(*t*) as our measure of coherence; similar effects were observed using the Pearson correlation.

As shown in Figure 5, the coherence between two regions increased as sodium (Fig. 5a) and calcium (Fig. 5b) were hyperpolarized, providing greater coupling between regions without varying the coupling constant *c*. Similarly, as sodium and calcium potentials were depolarized, inter-regional coherence decreased, leading to less synchrony between regions. We observed a similar pattern in potassium (Fig. 5c), where the depolarization of the potassium ion gradient leads to greater coupling (due to significantly slower oscillations as the refractory period lengthens). The effects of potassium are noticeably smaller than those of sodium and calcium, likely due to a smoother gradient with the lack of the torus bifurcation.

**Figure 5:**
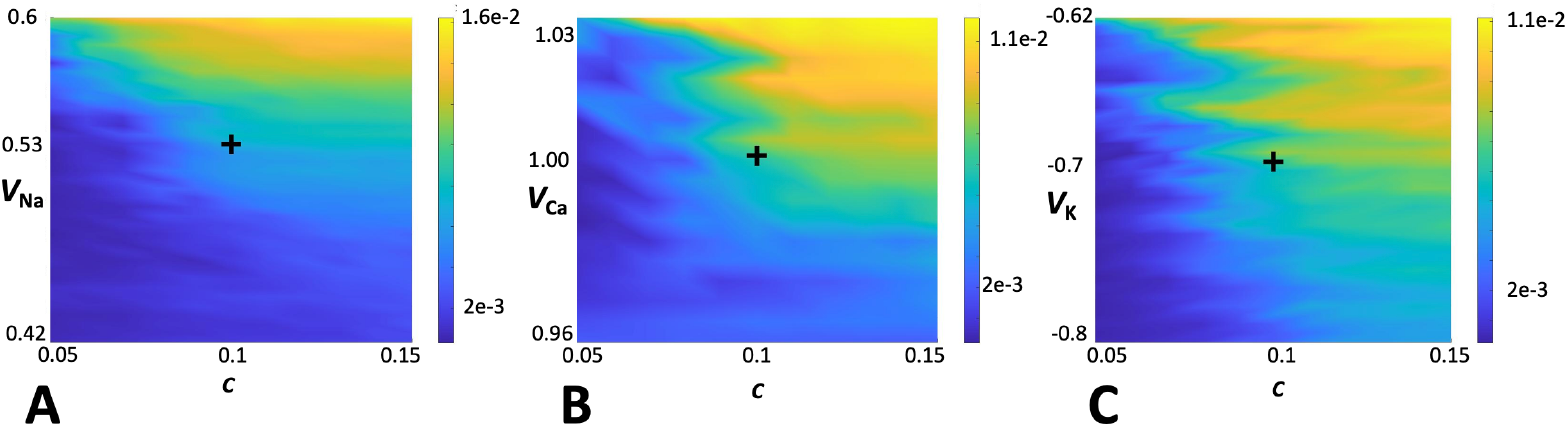
Coherence is poised at a critical point of both ion reversal potentials and inter-region coupling. Coherence (measured as covariance) between coupled neural masses as a function of coupling constant *c* and ion gradients. The color bars show the covariance of the neural time series. All center points have a *c* = 0.1. A. Coherence between two neural masses as a function of *V*_Na_, where baseline sodium potential is shown at the cross (*V*_Na_ = 0.53). B. Coherence between two neural masses as a function of *V*_Ca_, where baseline calcium potential is shown at the cross (*V*_Ca_ = 1.0). C. Coherence between two neural masses as a function of *V*_K_, where baseline potassium potential is shown at the cross (*V*_K_ = −0.7).

The crosses in Figure 5 show the location of the baseline coupling between neuronal masses. These are located at the intersection of the physiological ion reversal potentials and the marginally stable coupling coefficient. We observe that these are poised at critical points, where a change in either ion gradient or coupling coefficient induces a sharp change in coherence between regions. As sodium and calcium reversal potentials are depolarized, the coupling coefficient must increase to compensate (i.e., coupling at the same level produces less synchrony when the ion gradient is depleted). Conversely, the hyperpolarization of sodium and calcium allowed for a smaller coupling constant to achieve the same synchrony, indicating greater coupling between regions induced by the neuronal dynamics stabilized by increased ion gradients. Changes in potassium exhibited similar, though smaller, deviations from the critical point as well. Hyperpolarization of potassium required an increase in the coupling constant, while depolarization of potassium allowed a lower coupling constant to achieve the same synchrony.

## 5. Discussion

### 5.1. Physiological relevance of bifurcations in ion gradients

In this work, we have demonstrated the existence and location of torus and flip bifurcations in the sodium ion gradient of the Larter-Breakspear model. While the model is well established [12, 13, 24], the manipulation of ion gradients has not been examined in prior literature. Physiologically, sodium serves to depolarize the neuron membrane when an action potential is fired [1, 2]. In this way, sodium governs the amplitude of the action potential, which in turn determines the length of the refractory period (i.e., higher amplitude requires a longer refractory period to recover). The torus bifurcation in the sodium gradient occurs depolarized relative to the baseline reversal potential. As the torus is crossed, we observe a loss of peaks in the membrane potential *V*, meaning depolarization has become so severe that the neurons are incapable of firing normal action potentials. Instead, they settle on small amplitude noisy oscillations that no longer resemble functional neuronal activity. This is consistent with single neuron studies that show under severe metabolic constraints, the loss of sodium reversal potential leads to a lack of neuronal activity [2, 11]. Conversely, the flip bifurcation in the sodium gradient occurs hyperpolarized relative to the baseline reversal potential. Crossing the flip bifurcation causes a significant increase in the amplitude of peaks in *V*, which in turn leads to a prolonged refractory period.

We have also shown the existence and location of torus and flip bifurcations in the calcium ion gradient. Under homeostatic conditions, calcium serves to aid sodium in the depolarization needed to fire an action potential, while also serving as an intracellular signaling molecule [11, 25]. Crossing the torus bifurcation leads to a significant depolarization in the calcium gradient, causing a loss of spikes similar to that observed in sodium depolarization. Crossing the flip bifurcation, however, causes only a very modest increase refractory time. While calcium is required to fire an action potential, it does not contribute to the amplitude in the same way as sodium [8], and so hyperpolarization with calcium does not exhibit as marked an effect on the refractory period after spiking.

In the case of potassium, we have shown the existence and location of only a flip bifurcation in the ion gradient. Mathematically, this is due to the unique modeling of potassium as a third differential equation in the Larter-Breakspear system. Physiologically, this is done to capture the importance of potassium as the main ion responsible for repolarization after an action potential is fired [9, 12]. When flip bifurcation is crossed and the potassium reversal potential is depolarized, there is a significant increase in the refractory period, reflecting the additional work that must be done to restore the membrane potential without a robust potassium gradient. There is no torus bifurcation in the potassium ion gradient as it is hyperpolarized. Physiologically, this is manifested as an increase in oscillatory frequency with no significant change in the shape of the peaks. This increased frequency is due to the presence of additional potassium, allowing for more rapid repolarization after firing an action potential. Since sodium and calcium are unchanged, we observe no alteration in the main peak shape.

The existence of codimension 1 (and occasionally codimension 2) bifurcations has been previously reported in both neural mass models and networks of spiking neurons [18, 19, 21, 29, 30]. Significantly, in networks of coupled neurons (rather than neural masses), chaotic quasi-periodic behavior such as that observed in actual neurons was observed to exist in a space lying between torus and flip bifurcations [21, 26]. This is similar to the torus and flip bifurcations we propose as the physiologically relevant boundaries in the Larter-Breakspear model, giving us greater confidence that these are, indeed, reflecting underlying neurobiological constraints.

### 5.2. Effects of ion gradient variability on inter-region coupling

Having derived these bifurcations, we then show that they are useful in examining the dynamics of coupled neural masses. Synchrony between brain regions is required for neurological function and deteriorates when metabolic constraints become too severe [4, 5]. At the neuronal level, these metabolic constraints cause the deterioration of the ion gradients needed for neural activity [2, 11]. We show that the depolarization of sodium and calcium ion potentials causes decreased coherence between coupled regions, effectively reducing their synchrony as though they were more loosely coupled. Similarly, hyperpolarization of sodium and calcium gradients produced greater inter-regional coherence, thereby promoting synchrony. We also demonstrate that depolarizing the potassium reversal potential increases inter-region coherence, while the hyper-polarization of potassium decreases coherence (due to increased oscillation frequency). Taken together, these results highlight the importance of stabilizing ion gradients to promote network synchrony, which has been shown in recent metabolic studies as well [5, 6].

We demonstrate that the coupled regions exist at a critical point between the ion gradients and coupling constant, with a small variation in either causing a large shift in the synchrony between regions. Operation at a critical point has been demonstrated to be a key feature of neuronal dynamics [31]. Brain regions exist in marginally stable coupling, allowing for individual oscillations within regions while retaining periodic synchronization across regions [13, 24]. Operating at a critical point may allow neuronal systems greater flexibility to respond most effectively to physiological perturbations [13, 31]. When depolarizing the sodium and calcium ion gradients (or hyperpolarizing the potassium ion gradient), we showed a perturbation from the critical point, requiring a greater coupling constant to achieve a similar level of synchrony. Physiologically, the new coupling constant will be difficult to achieve, and so the loss of an ion gradient will correspond to a decrease in synchrony between regions, as has been observed in metabolically constrained networks [4, 5].

### 5.3. Limitations and future directions

The primary limitation of this work is the lack of purely analytical derivation of the bifurcation points. While analytical discovery of bifurcations is preferable, the complexity of the Larter-Breakspear model is such that the derivation of exact solutions would be extremely difficult. The numerical analysis of bifurcations is also well-established characterizing chaotic dynamics [16, 27, 20], and has become a standard means of analyzing complex systems.

Perhaps the most useful extension of this work would be the analysis of bifurcations when there is time-delayed coupling between regions. There has already been work [24, 32] demonstrating that time-delays and the refractory period in neural mass models can model the propagation of electrical waves throughout the brain. There has also been prior work on how the introduction of time delays in chaotic systems [18], including neural mass models [33], can introduce novel bifurcation points.

## 6. Conclusion

We present here a detailed characterization of the ion gradients in the Larter-Breakspear model, an aspect of the model that has not been explored in prior works. We show the existence and location of torus and flip bifurcations in the sodium and calcium ion gradients, and a single flip bifurcation in the potassium ion gradient. These bifurcations correspond to physiologically relevant limits, with depolarization or hyperpolarization beyond these bifurcations causing significant alteration in the shape and frequency of neural mass spiking patterns. We also demonstrate that in a system of coupled neural masses, depolarization of the sodium and calcium ion gradients leads to decreased inter-region coherence, while hyperpolarization increases inter-region coherence. These results further emphasize the role of metabolism in maintaining network stability within the brain, and the model parameters explored here can be used in future modeling to build multi-scale simulations that probe the functional correlates of metabolic constraints.

## 7. Acknowledgments

The research presented here was funded by the W. M. Keck Foundation (to LRMP), the White House Brain Research Through Advancing Innovative Technologies (BRAIN) Initiative (NSFNCS-FR 1926781, also to LRMP), and the Baszucki Brain Research Fund (also to LRMP). Anthony Chesebro was also supported by the NIHGM MSTP Training Award T32-GM008444. Corey Weistuch also acknowledges the Marie-Josée Kravis Fellowship in Quantitative Biology. We would like to thank Dr. Okito Yamashita for his prompt and helpful input during our early development of the Larter-Breakspear model code.

## 8. CRediT statement

**Anthony Chesebro**: Conceptualization, Formal analysis, Software, Validation, Visualization, Writing-original draft, reviewing and editing. **Lilianne Mujica-Parodi**: Conceptualization, Funding acquisition, Writing-reviewing and editing. **Corey Weistuch**: Conceptualization, Formal analysis, Visualization, Writing-original draft, reviewing and editing, Supervision.

## 9. Code Availability

All code used to create the models and generate the results presented in this paper is available at https://github.com/agchesebro/larter-breakspear-bifurcation. Bifurcation analysis was performed with the MatCont toolbox [27], available to download at https://sourceforge.net/projects/matcont/. Implementation of the analyses presented here that were performed in MatCont are available at the model link above.

## Appendix A. Notes on selection of *δ*_*V*,*Z*_

The range of the excitatory and inhibitory membrane threshold (*δ*_*V*,*Z*_) has been the subject of much prior work in the Larter-Breakspear model [12, 13, 14]. The range of the parameter is fairly narrow; *δ*_*V*,*Z*_ < 0.59 fails to produce oscillations and instead results in a model that immediately converges to an equilibrium point, while *δ*_*V*,*Z*_ > 0.7 produces non-physiological dynamics [14].

**Table B.2:** Critical points in the ion gradients.

| Parameter | Limit Point | Neutral Saddle Equilibrium |
| --- | --- | --- |
| $V_{Na}$ | -1.312 | 2.432 |
| $V_{Ca}$ | -1.196 | 1.552 |
| $V_K$ | -1.242 | 0.321 |

We set *δ*_*V*,*Z*_ = 0.66 in this work as it achieves consistently useful results. Selecting a different value may shift the exact thresholds we discuss here, but remaining within its operational range does not change quality of the ion gradient landscapes we discuss.

## Appendix B. Other critical points in the ion gradients

For all three ions, we found that in additional to the Hopf points considered above, there also exist a neutral saddle equilibrium point above and a limit point below the Hopf points. These are presented in Table 2 for completeness; however, we note that they are so far beyond the physiologically relevant limits discussed in this text that they are more of theoretical interest than practical value. We also noted that in most cases they produce degenerate orbits (i.e., the values are too far removed from equilibrium to even induce cycling and tend to converge on a nonsense value). The one exception to this is the limit point of potassium which, although physiologically improbable, still produces discernible periodic orbits. This is most likely due to the lack of a torus bifurcation in the potassium gradient.

